# De novo identification, differential analysis and functional annotation of SNPs from RNA-seq data in non-model species

**DOI:** 10.1101/035238

**Authors:** Hélène Lopez Maestre, Lilia Brinza, Camille Marchet, Janice Kielbassa, Sylvère Bastien, Mathilde Boutigny, David Monnin, Adil El Filali, Claudia Marcia Carareto, Cristina Vieira, Franck Picard, Natacha Kremer, Fabrice Vavre, Marie-France Sagot, Vincent Lacroix

## Abstract

SNPs (Single Nucleotide Polymorphisms) are genetic markers whose precise identification is a prerequisite for association studies. Methods to identify them are currently well developed for model species, but rely on the availability of a (good) reference genome, and therefore cannot be applied to non-model species. They are also mostly tailored for whole genome (re-)sequencing experiments, whereas in many cases, transcriptome sequencing can be used as a cheaper alternative which already enables to identify SNPs located in transcribed regions. In this paper, we propose a method that identifies, quantifies and annotates SNPs without any reference genome, using RNA-seq data only. Individuals can be pooled prior to sequencing, if not enough material is available for sequencing from one individual. Using human RNA-seq data, we first compared the performance of our method with Gatk, a well established method that requires a reference genome. We showed that both methods predict SNPs with similar accuracy. We then validated experimentally the predictions of our method using RNA-seq data from two non-model species. The method can be used for any species to annotate SNPs and predict their impact on proteins. We further enable to test for the association of the identified SNPs with a phenotype of interest.

## 1 Introduction

Understanding the genetic basis of complex phenotypes remains a central question in biology. A classical approach consists in genotyping a large number of individuals in a population based on a pre-specified catalog of variants, and in associating their genotypes to the studied phenotype. This type of approach can be applied to many loci at once, or even genome wide, through what has been called genome wide association studies (GWAS). These methods have been successfully adopted for human and other model species. However, the total cost of GWAS remains very high, and the current framework cannot be applied to non-model species for which genomic resources are sparsely or not available. The recent progress in sequencing technologies together with the recent developments in assembly algorithms are largely changing this view. It can now be envisioned to search for variants associated with a phenotype using NGS data only, without relying on pre-existing genomic resources (that have potential limitations). A possible procedure, applicable to model or non-model species, consists in: (1) sequencing the genome; (2) assembling it; (3) identifying the SNPs; (4) genotyping individuals; and (5) associating genotypes with phenotypes. However, such a procedure remains costly and still presents the classical problems of sequential pipelines, namely the potential to accumulate experimental and computational errors at each step.

If the purpose of the study is to identify the variants related to a phenotype, the procedure can be simplified in many ways. First, SNPs can be called *de novo* from the reads, without separating the steps of assembly and SNP calling. Second, cost effective methods like exome or transcriptome sequencing may be adopted as the full genome is not always necessary. Third, pooling individuals may be an attractive option if genotyping is not required. These options have been explored individually and give promising results. *De novo* assembly of SNPs is now computationally possible [Iqbal et al., 2012, Uricaru et al., 2015, Leggett et al., 2013]. The clear advantage is that it can be applied to non-model species, where no reference genome is available. Even in the case where a reference genome is available, these methods still give good results compared to mapping-based approaches, compensating their lower sensitivity by an ability to call more variants in repeated regions. Transcriptome sequencing is already used in several projects, both in the context of model species [Piskol et al., 2013] and nonmodel species [Romiguier et al., 2014]. In both cases, it was shown that the SNP calling methods could be tailored to have a good precision, meaning that most of the reported SNPs are true SNPs. However, their recall (i.e. capacity to exhaustively report all SNPs) remains to be clearly determined. Clearly, only SNPs from transcribed regions can be targeted, but they arguably correspond to those with a more direct functional impact. Using RNA-seq technology largely reduces the cost of the experiment, and the obtained data concurrently mirror gene expression, the most basic molecular phenotype. RNA-seq experiments may also provide very high depth at specific loci and therefore allow to discover infrequent alleles in highly expressed genes. Finally, pooling samples is already extensively used in DNA-seq (sometimes termed Pool-seq) [Schlötterer et al., 2014]. The main advantage of this method is that it clearly decreases costs, as library preparation for bar-coding is nowadays roughly the same price as sequencing. The drawback is that genotypes cannot be derived anymore. Instead, we have access to the allele frequency in the population, a result known as the allelotype.

In this work, we present a method for the de novo identification, differential analysis and annotation of variants from RNAseq data in non-model species. It takes as input RNA-seq reads from at least 2 conditions (e.g., the modalities of the phenotype) with at least 2 replicates each, and outputs variants associated with the condition. The method does not require any reference genome, nor a database of SNPs. It can therefore be applied to any species for a very reasonable cost. We first evaluated our method using RNA-seq data from the human Geuvadis project [Lappalainen et al., 2013b]. The great advantage of this dataset is that SNPs are well annotated, since the selected individuals were initially included in the 1000 genomes project [Abecasis et al., 2012]. This enables to clarify what is the precision and recall of our method, and how it compares to widely used methods like Gatk, which require a reference genome.

We then applied our method in the context of non-model species. First we focused on *Asobara tabida*, an hymenoptera that exhibits contrasted phenotypes of dependance to its symbiont. Using RNA-seq data from two extreme modalities of the phenotype, we were able to establish a catalog of SNPs, stratify them by functional impact, and assess which SNPs had a significant change of allele frequency across modalities. We further selected cases for experimental validation, and were able to confirm that the SNPs were indeed condition specific. Then we applied our method on two recently diverged *Drosophila* species, *D. arizonae* and *D. mojavensis*. These species can still produce hybrids that are sterile. In this case, our method identifies differences of 1 nt, which are not SNPs but divergences. On this system also, we were able to validate experimentally that the loci we identify were truly divergent.

We outline that, even though the case studies presented in this paper include two replicates, the method can be applied to any number of replicates. Larger cohorts can be helpful to narrow down the list of SNPs likely to be really causal for the phenotype. Our key contribution is that we are able to produce a list of SNPs stratified by functional impact, and ranked by difference of expressed allele frequency across conditions. This list can be further mined for candidates to follow up experimentally.

All the methods presented in this paper are implemented in software that are freely available at http://kissplice.prabi.fr/TWAS. In particular, the statistical procedure that we developed is available through an R package, KISSDE, which is of general interest for researchers who have obtained read counts for pairs of variants in a set of conditions and wish to test if these counts reflect the specificity of the variant in a particular condition.

## 2 MATERIALS AND METHODS

### 2.1 Overview

We present here a collection of methods which can be used together to produce, from RNA-seq data alone, a list of condition-specific SNPs, stratified by their predicted impact on the protein. Figure 1 summarises the different steps.

**Figure 1:**
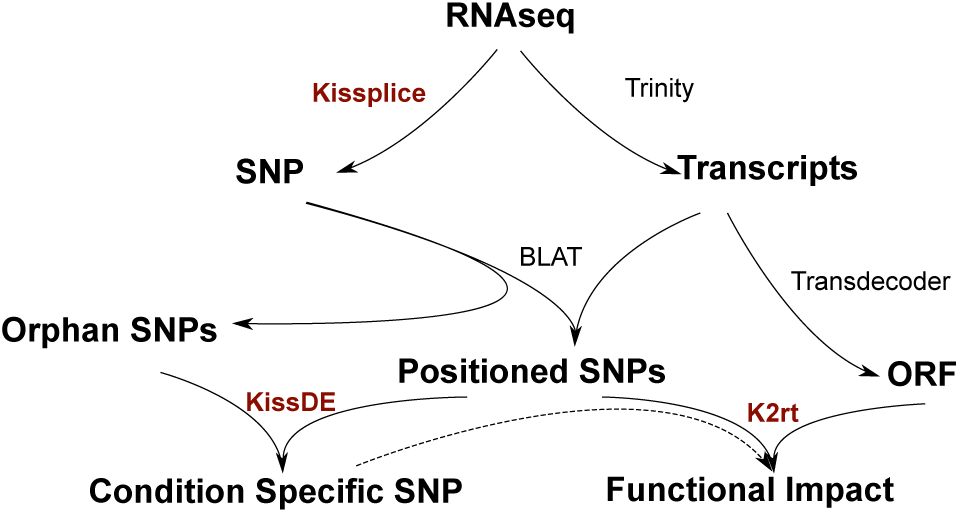
With fasta/fastq input from an RNA-seq experiment, SNPs are found by Kis-Splice without using a reference. As KisSplice provides only a local context around the SNPs, a reference can be built with Trinity, and SNPs can be positioned on whole transcripts. Some SNPs that do not map on the transcripts of Trinity, called orphan SNPs, are harder to study but can still be of interest. We propose a statistical method, called KissDE to find condition-specific SNPs (even if they are not positioned) out of all SNPs found. Finally, we can also predict a functional impact for the positioned SNPs, and intersect these results with condition-specific SNPs using our package KisSplice2-RefTranscriptome (K2rt).

Trinity, Transdecoder and Blat are third-party software. KisSplice was published recently [Sacomoto et al., 2012], kissDE and KisSplice2RefTranscriptome (K2rt) are methods we introduce in this paper.

### 2.2 De novo identification of SNPs

KisSplice [Sacomoto et al., 2012] is a software initially designed to find alternative splicing events (AS) from RNA-seq data, but which also outputs indels and SNPs. We present here its functionality for SNP detection. The key concept, initially introduced in [Peterlongo et al., 2010] and later used in [Iqbal et al., 2012, Uricaru et al., 2015] is that a SNP corresponds to a recognisable pattern, called a *bubble*, in a de Bruijn graph (DBG) built from the reads. De Bruijn graphs are widely used data structures in de novo assembly [Pevzner et al., 2001, Zerbino and Birney, 2008, Grabherr et al., 2011], as they are well tailored for large amounts of short reads. In our case, DBGs are especially appealing because they model explicitly each nucleotide, a required feature to capture SNPs. The nodes of the graph are words of length k, called k-mers. There is an edge between two nodes if the suffix of length *k* − 1 of the first k-mer is identical to the prefix of length *k* − 1 of the second k-mer. The DBG that is built from two alleles of a locus will therefore correspond to a pair of vertex-disjoint paths in the graph, which form the bubble. Unlike AS events and indels, bubbles generated by SNPs have two paths of equal length (Figure 2). Notice that additionally, linear paths of the DBG can be further compressed in a single node without loss of information (Figure 2).

**Figure 2:**
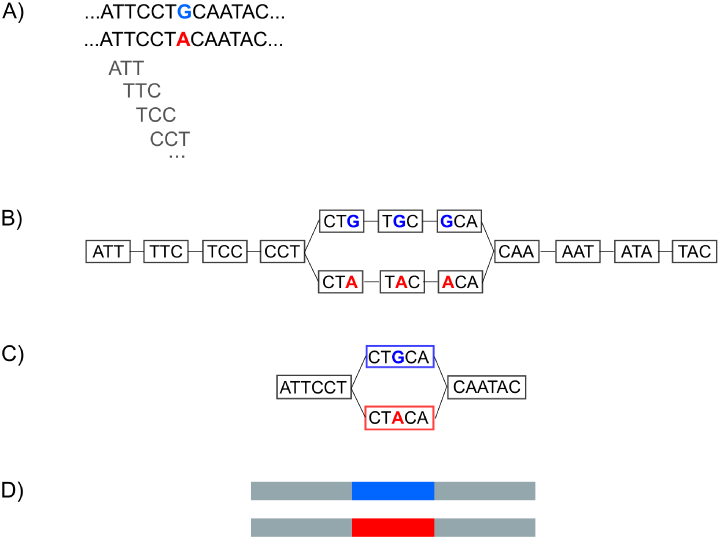
A) A SNP present in two alleles in the data. B) The de Bruijn Graph derived from the data. For the sake of simplicity of exposition, we draw here with *k*=3. In practice, *k*=41. C) A compressed de Bruijn Graph can be obtained by merging nodes with a single outgoing edge with nodes with a single incoming edge. This compression step is lossless. D) The two paths in the compressed de Bruijn Graph correspond to the two alleles of the SNP.

In the special case where there are two SNPs located less than *k* nt apart on the genome, they will be reported in the same bubble (Supplementary Figure 1). In the case where the two SNPs are perfectly linked, a single bubble is reported. If they are partially linked, each haplotype will correspond to a path, and KisSplice will report all pairs of paths. In this case, the number of bubbles does not correspond to the number of SNPs, but to the number of pairs of observed haplotypes. Supplementary Figure 2 illustrates the case of 2 SNPs and 4 haplotypes.

KisSplice consists in essentially 3 steps: (i) building the DBG from the RNA-seq reads; (ii) enumerating all bubbles in this graph; and (iii) mapping the reads to each path of each bubble to quantify the frequency of each variant. Particular attention was paid to both the memory [Chikhi and Rizk, 2013, Salikhov et al., 2014] and time [Sacomoto et al., 2014] requirements of the pipeline. KisSplice was able to process 200M reads of 2*75nt in 20 hours, with less than 16G of RAM.

### 2.3 Filtering out sequencing errors and inexact repeats

SNPs correspond to bubbles in the de Bruijn graph derived from the reads. However, not all bubbles in the DBG correspond to SNPs. Essentially two types of false positives can be found: sequencing errors and inexact repeats. RNA editing sites may also be mistaken for SNPs but in practice, these correspond to a few cases only, that we discuss in the Results section.

**Sequencing errors** may generate bubbles in the DBG. A distinctive feature that helps to discriminate them from true variants is that one path of the bubble is expected to be poorly covered. In practice, a common way to filter out sequencing errors when dealing with DNA-seq data is to remove all rare k-mers (seen less than a given number of times) prior to the DBG construction. This simple strategy, implemented for instance in DiscoSNP, is however not sufficient when dealing with RNA-seq data. Since the coverage depends on gene expression, it is therefore very unequal across genes, and the cut-off should be adapted to each gene. To account for this constraint, we introduced a relative cut-off, which enables to remove edges in the DBG that are supported by less than a percentage of all counts outgoing from (or incoming to) the same node. This enables to remove sequencing errors even in highly expressed genes (Figure 3). Clearly, the drawback of these cut-off strategies is that rare variants will be filtered out because they will be mistaken for sequencing errors. Our ability to detect rare variants is therefore limited by this critical parameter. In this study, we set the cut-off to 5%.

**Figure 3:**
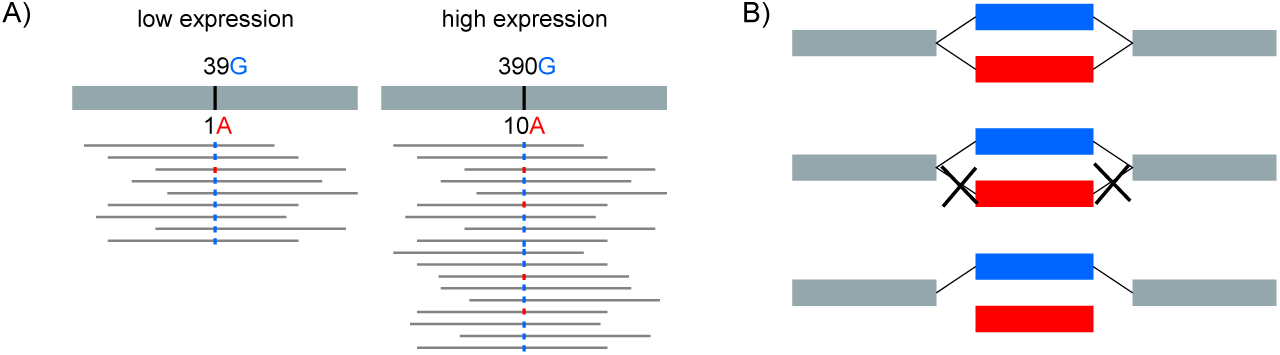
Sequencing errors and rare variants generate bubbles in DBGs with very unbalanced path coverage. (A) For ease of exposition of the concept, we represent here the reads mapping to a reference genome. Applying an absolute cutoff would remove the sequencing error for a poorly expressed gene, but not for a highly expressed gene. B) Applying a relative cutoff of 5% in the DBG removes one or two edges from the red path and hence prevents this bubble from being found.

**Inexact genomic repeats** may also generate bubbles in the DBG (Figure 4). This is the case for instance for recently diverged paralogs which still share a lot of sequence similarity and hence may differ locally by one nucleotide flanked by *k* conserved nucleotides. This is also the case for other types of repeats, including inexact tandem repeats or transposable elements which may be present in the UTRs and introns of genes. In principle, introns are not present in RNA-seq data, but in practice, whatever the protocol used to filter out pre-mRNA, a proportion of at least 5% remains [Tilgner et al., 2012].

**Figure 4:**
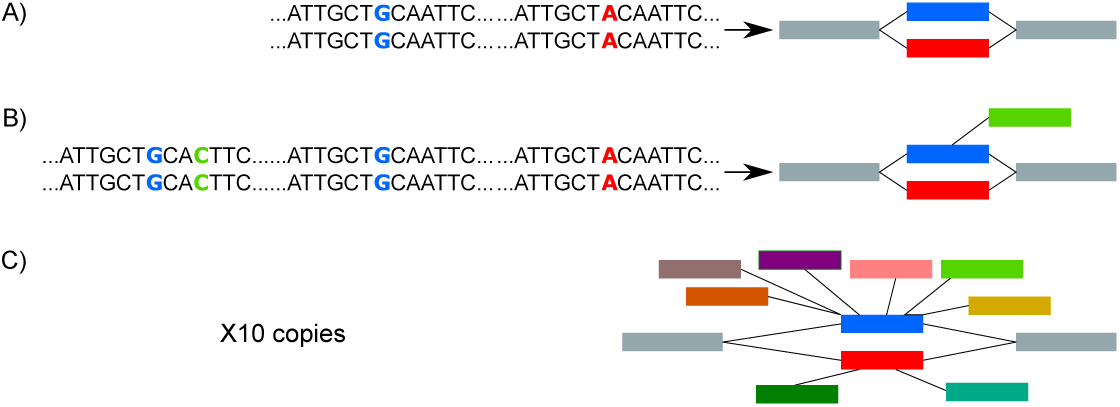
Two inexact repeats give rise to a pattern in the DBG that resembles a SNP (A). Very often, repeats are present in more than 2 copies (B) and therefore generate branching bubbles. Bubbles with more than 5 branches (C) are filtered out.

The question of discriminating SNPs from inexact repeats has already been addressed in the literature in the case of unpooled data. Romiguier et al. [Romiguier et al., 2014] propose to use the idea that loci corresponding to recently diverged paralogs should present an excess of heterozygous sites. This idea cannot be employed in our case since we want our method to be able to deal with pooled data, where we cannot genotype individuals. Besides, it does not apply to cases where only a portion of the transcript contains repeats, which happens often, when some pre-mRNA is still present in the samples.

Repeats present in a large number of copies (like transposable elements, or large families of paralog genes) generate a large number of bubbles which are false positives. However, these bubbles have a specific feature that we can use to discriminate them from the others: they are branching (Figure 4). The more (inexact) copies in the repeat family, the higher the number of branches in each bubble. In order to filter them out, we introduced a parameter *b*, which corresponds to the maximum number of branches allowed. If one path of the bubble has more than *b* branches, then the bubble is filtered out. In practice, we set this parameter to 5, which appeared to be a good trade-off between recall and precision.

Repeats present in a small number of copies are not filtered out by our criterion. Nevertheless, most of them are actually filtered out at the next step of the pipeline, when we test for the enrichment of one variant in one condition (as described in the Statistical analysis section). The ones that are not filtered out at this step correspond to paralogous genes, where one copy is more expressed in the first condition and the second copy is more expressed in the other condition. Although these are not SNPs, we can argue that they are still relevant candidates for an association study aiming at proposing causes for the difference of phenotype.

### 2.4 Predicting the functional impact of SNPs

KisSplice predicts SNPs, but outputs only a very local context around the SNP. In order to predict its functional impact, we need to place the SNP in a larger genomic context. For this, we relied on a widely used global transcriptome assembler: Trinity [Grabherr et al., 2011], which takes as input RNA-seq reads and outputs contigs that correspond to either full-length transcripts (if the expression level of the transcript is sufficient) or to fragments of transcripts. The results of KisSplice were aligned onto the transcripts predicted by Trinity using Blat [Kent, 2002]. Concurrently, we searched for coding potential in the transcripts using TransDecoder. Once we had the location of the SNP within the transcript and the location of the open reading frame (ORF), we could assess if the SNP was located within the CDS or not, and if so, if it was a synonymous or non synonymous SNP. In the case where no ORF was predicted for the transcript, we concluded that the SNP was within a non coding region. In practice, this can correspond to a non coding RNA, a UTR or an intron. Prediction of the functional impact of a SNP was included in a Python package, called KisSplice2RefTranscrip-Tome (k2rt), which takes as input a set of predicted ORFs (bed format), the output of KisSplice (fasta format), and a mapping of the results of KisSplice to the transcripts (psl format). Importantly, Trinity, TransDecoder and Blat are third party software which can be replaced by others, provided the exchange formats are respected (bed and psl).

In the case where a SNP mapped to several Trinity transcripts, we reported the functional impact of the SNP in each transcript. This happened in particular when a SNP was located in a constitutive exon of a gene that gave rise to multiple alternative transcripts through alternative splicing.

In the case where a SNP mapped to no transcript, then it could not be treated by k2rt and it was filtered out. Those SNPs were called orphan SNPs. They were mostly located in poorly expressed genes and/or highly repeated regions. Indeed, repeated regions are notoriously difficult to assemble. When repeated regions are located within genes, they may either generate chimeric transcripts in the assembly if the assembler is too permissive, or a series of truncated short contigs if the assembler is too conservative. By default, Trinity does not output contigs shorter than 200 nucleotides. Because these contigs are highly enriched in repeats and poorly expressed genes, it explains the origin of the majority of our orphan SNPs.

As mentioned in the model section, the number of bubbles does not always correspond to the number of SNPs. In the case of SNPs located less than *k* nucleotides apart, the number of bubbles corresponds to the number of pairs of haplotypes out of the total number of haplotypes. The same SNP may therefore be present in multiple bubbles. When mapping the bubbles to a reference transcriptome, it is possible to remove this redundancy and count the true number of SNPs. Indeed, if two bubbles map to the same transcript at the same location, then it means that they refer to the same SNP, and we count it only once.

The software versions that we used were: Trinity r20140717, TransDecoder v2.0.1, BlatSuite36, KisSplice v2.4, KisSplice2RefTranscriptome v1.0.

All were used with default parameters, except Blat where we set --minIdentity=80. We also set the minimum query coverage to 80% in k2rt. Changing both the minimum identity and the minimum query coverage from 70% to 90% only marginally affected our results.

A critical parameter in de novo assembly is the *k*-mer size. In Trinity, this value is set to 25 and cannot be modified. In KisSplice the default value is 41 as we found it is a good compromise between recall and precision. We also tested 25 and this resulted in an increase of 10% in recall but a decrease of 10% in precision. For advanced users interested in obtaining a more exhaustive list of candidates (hence optimising recall), we recommend to decrease the value of *k* in KisSplice.

### 2.5 Statistical analysis

#### 2.5.1 Testing the association between a variant and a condition

Given the number of SNPs (*n*) and the number of replicates (*m*), our dataset is a count matrix of size 2*n* × *m*, with 2 lines corresponding to 1 SNP (upper and lower path representing the two different alleles with one nucleotide differing between both paths). Before comparing the allele read counts from different libraries, the count data were normalised by library sizes as proposed in the DESeq package [Anders, 2010]. This software has been shown to be the most efficient according to a recent normalisation comparison study [Dillies et al., 2013]. Some events were then filtered out based on their counts: if global counts (for all replicates and all conditions) for both variants were too low (< 10), we considered that we did not have enough power to conclude on this event and we did not test it.

Our statistical analysis adopted the framework of count regression with Negative Binomial distribution as in standard RNA-seq analysis. We considered a 2-way design with interaction, with *alleles* and *experimental conditions* as main effects. Following the Generalized Linear Model framework, the expected intensity of the signal was denoted by *λ_ijk_* and was decomposed such that:

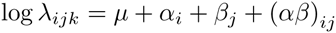

where *μ* is the local mean expression of the transcript that contains the SNP, *α_i_* the effect of allele *i* on the expression, *β_j_* the contribution of condition *j* to the total expression, and *(αβ)_ij_* the interaction term. The target hypothesis is *H*_0_: {(*αβ*)*_ij_*· = 0} (*n*o interaction between allele and condition) that is tested using a Likelihood Ratio Test [Bullard et al., 2010] with a 5% false discovery rate (FDR) to account for multiple testing.

Due to numerical instabilities associated with the estimation of Negative Binomial parameters, we adopted a model selection approach to determine the best model to handle the over-dispersion parameter (*ϕ*). Our strategy was first to estimate a model without over-dispersion using the Glmnet package (model 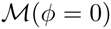. We then considered two different estimation methods for the parameter *ϕ*, namely a global estimation approach using the package aod (model 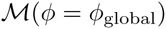)), and a SNP-specific parameter using the DSS package (model 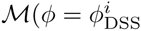.We finally used a BIC to choose the best model out of the three. Pseudo-counts (*ie*, systematic random allocation of ones) were considered for SNPs showing many zeros to avoid singular hessian matrices while fitting the generalised linear model.

#### 2.5.2 Quantifying the magnitude of the effect

When a variant is found to be differentially represented in two populations, one remaining difficulty is to quantify the magnitude of this effect. Indeed, significant *(p <* 0.05) but weak effects are often detected, especially in RNA-seq data in which some genes are very highly expressed (and hence have very high read counts).

A natural measure for quantifying the magnitude of the effect would be the difference of allele frequencies between the two conditions. In practice, the true difference of allele frequencies is not known, and we estimated it using the RNA-seq counts. The precision of this estimation is discussed in the Results Section.

We denote by *f_e_* the estimation of the allele frequency based on RNA-seq counts:

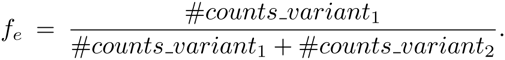

The value of *f_e_* was computed for each replicate of each condition. We then took the mean of these values for all replicates within each condition. Finally, we calculated the difference across conditions and obtained the magnitude of the effect: 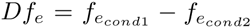. In the special case where the two variants had low counts (< 10) within one replicate, then *f_e_* was not calculated. Finally, if at least half of the replicates of one condition had low counts, *Df_e_* was not computed either. Overall, this prevented from over-interpreting large magnitudes obtained from low counts.

Our method is embedded and distributed in an R package, called KissDE which can take as input either the output file of KisSplice or any count matrix with two lines representing an event.

### 2.6 Methodology for testing and validating our approach

We first evaluated our method in human, because it is a model species, for which a reference genome is available and SNPs are well annotated. We then used our method on a non-model species: *Asobara tabida*, an hymenoptera that exhibits contrasted phenotypes and for which no reference genome is available. Finally, we applied our method on a different evolutionary timescale, working on two recently diverged *Drosophila* species, *D. mojavensis* and *D. arizonae*, for which a draft reference genome is available only for *D. mojavensis*.

#### 2.6.1 The Geuvadis dataset

Our method enables to find SNPs from RNA-seq data. In order to assess if the SNPs we find are correct, and if the list we output is exhaustive, we chose to test our method on RNA-seq data from the Geuvadis project. Indeed, the individuals whose transcriptome was sequenced in this project were already included in the 1000 genome project. Hence, their SNPs have been already well annotated. We downloaded fastq files from SRA (see Data access) and selected 10 Toscans and 10 Central Europeans. We sampled 10M reads for each individual and concatenated the fastq files in pools of 5 individuals.

*Definition of the set of true SNPs and their genotypes* We downloaded the vcf file from the 1000G webpage. For each SNP called in the 1000 Genomes project, we had at our disposal the genotype of each individual. We focused on the genotypes of the 20 individuals selected for our analysis. Whenever only one allele was represented in the 20 individuals, we filtered out this SNP, as it simply cannot be discovered based on these 20 individuals only.

Whenever one SNP was covered by less than 5 reads out of the total number of reads in the 20 individuals, we considered that the SNP was located in a too poorly expressed region and could not be discovered by RNA-seq. Other levels of poorly/medium/highly expressed regions are discussed in the Results section. The read coverage was computed using Samtools depth, on the .sam file obtained after mapping the reads with Star (v2.3.0) [Dobin et al., 2013].

##### *Prediction of SNPs with the mapping-based approach:* Gatk

In order to clarify if the performances of our method were on par with other methods, we chose to benchmark against Gatk, which is the most widely used method for variant calling in eukaryote samples.

We employed the Gatk Best Practices workflow for SNP and indel calling on RNA-seq ^1^ data which considers the following steps: (1) mapping to the reference with the Star aligner, 2-pass method ([Engström et al., 2013]) with the suggested parameters allowing to obtain the best sensitivity for variant call task, where during the second pass of Star a new reference index is created from the splice junction information determined during the first step alignment and a new alignment step is done with the new index reference; (2) adding read group information, sorting, marking duplicates and indexing, using Picard’s tools; (3) splitting reads into exon segments (removing Ns but maintaining grouping information) and hardclipping sequences overhanging into the intronic regions, using the SplitNCigarReads Gatk tool; (4) realigning indels and recalibrating Base quality; and finally (5) calling variant with HaplotypeCaller.

##### Comparison of mapping-based and assembly-based approaches

In order to be able to compare the results of KisSplice with the ones of Gatk, we needed to obtain genomic positions for all the SNPs assembled by KisSplice. For this purpose, we aligned each variant of each bubble to the reference genome using Star (2.3.0).

In the case where a variant mapped to several locations, we used the default behaviour of Star, which is to assign the variant to the location with the fewer number of mismatches. In case of ties, we kept all equally good locations and if at least one of the possible locations corresponded to an annotated SNP, then we considered that the prediction of KisSplice was correct.

#### 2.6.2 *Asobara tabida* lines, RNA sequencing and SNP verification

*Asobara tabida* (Hymenoptera: Braconidae) is a parasitoid species, which develops on *Drosophila* hosts. *A. tabida* is naturally infected by 3 strains of *Wolbachia*, among which one (*w*Atab3) is necessary for oogenesis completion ([Dedeine et al., 2001, Dedeine et al., 2004]). However, when *Wolbachia* are removed by antibiotic treatment, the degree of oogenetic defect exhibits genetic variation within populations [Kremer et al., 2010] We thus founded two lineages of *A. tabida* from a natural population (Sainte Foy-les-Lyon, France) based on their extreme phenotype after elimination of *Wolbachia*: the SFR2 lineage whose females do not produce any eggs and the SFR3 lineage whose females produce half the normal content of eggs. In both cases, dependence is complete as the eggs produced are sterile. These two lineages were founded by 3 females and were kept for 15 generations (three founders at each generation) before RNA extraction.

The experimental design for RNA-seq sequencing aimed at describing the transcriptomic changes associated with the presence / absence of *Wolbachia*, and the variations observed in the two *A. tabida* lineages exhibiting an extreme phenotype. For this purpose, cDNA libraries were constructed from infected and non-infected ovaries in these two lineages. Because these RNA-seq data were issued from two distinct lineages from a non-model species, we exploited this dataset to validate the method developed here and to discover biologically relevant SNPs, using libraries obtained from infected ovaries. The samples used for RNA extraction were young female (0–1 day old) ovaries dissected in a drop of A-buffer (2 replicates of 30 ovaries per lineage). RNA was extracted as described in [Kremer et al., 2009]. These RNA extracts were used to generate corresponding cDNA libraries, following the recommendations given by the manufacturer of the SMARTer PCR cDNA synthesis and BD Advantage 2 PCR kits (Clontech). These cDNA libraries were then purified with the Qiaquick kit (Qiagen) and their quality checked. Sequencing of cDNA was performed by the Genoscope (Evry), on an Illumina GA-IIx instrument, to obtain 1x75bp reads. These data were trimmed using the ShortRead package with default parameters and then used as input of the pipeline defined in Figure 1.

Based on these results, 27 SNPS were chosen for verification. For each SNP, primers were designed on the corresponding transcript to amplify the surrounding genomic region. PCRs were performed from an aliquot of the purified cDNA libraries. The reaction was performed in a total volume of 25 L, and the mixture consisted in 2.5 L of 5X green DreamTaq mastermix, 200 nM of dNTP, forward and reverse primers (see Supplementary Table 1 for primer sequences), and 5U of DreamTaq DNA polymerase (ThermoFisher). PCR amplification was performed on a Tetrad thermocycler (Biorad) as follows: 2 min at 94C, 35 times (30 s at 94C, 30 s at 58C, 30s at 72C), and 10 min at 72C. The PCR products were sequenced using the Sanger method from forward and reverse primers by the Biofidal company. The sequences were aligned and their respective chromatograms analysed by the CLC Main workbench.

#### 2.6.3 *Drosophila* strains, RNA sequencing and SNP verification

*D. mojavensis* and *D. arizonae* are two Drosophila species that are endemic of the arid southwestern United States and Mexico. These species diverged recently (less than 1 MYA [Matzkin, 2004, Reed et al., 2007]. In the laboratory, hybridisation of these two species is possible while in nature it does not occur (or is very rare). The ovarian transcriptome of these two species (and their reciprocal crosses) was sequenced to investigate the first step of hybrid incompatibility and look for deregulated genes in hybrids. In this paper, we did not study the transcriptomes of the hybrids, we only used the transcriptomes of the parents to test for the validity of our pipeline at a different evolutionary scale. The sequenced strains were *Drosophila mojavensis* from the Anza Borrego Desert, CA (stock number: 15081-1352.01) and *Drosophila arizonae*, from Metztitlan - Hidalgo, Mexico (stock number: 15081-1271.17), both obtained in the US San Diego Drosophila Stock Center. Virgin female flies were collected after hatching and isolated until they reached ten days. The RNA was extracted from a pool of 30 ovaries of 10-days-old flies for each line. The extractions were performed using the RNeasy kit (Qiagen) and samples were then treated with DNase (DNA-free Kit, Ambion) and stored at −80C. The samples were quantified by fluorescence in the Bioanalyser 2100 (Agilent), according to pre-established criteria by the sequencing platform. For each line, the extracted RNA was divided into two parts in order to generate two cDNA libraries (two replicates per condition). RNA was sequenced by Illumina Technology, in the IlluminaHiseq 2000. We sequenced 2x51bp paired-end reads and the medium size of the inserts was 300bp. We used UrQt [Modolo and Lerat, 2015] with the default parameters to remove the low quality bases and the polyA tail from the dataset before running the pipeline described in Figure 1. The protocol for SNP verification is identical to the one used for *Asobara tabida* (see Supplementary Table 2 for primer sequences).

### 2.7 Data access

The human data used in this study can be found through the ArrayExpress database (http://www.ebi.ac.uk/arrayexpress/) under the accession number E-GEOD-29342 and we used the individuals named NA20808, NA20809, NA20810, NA20811, NA20812, NA20813, NA20814, NA20815, NA20819, NA20826, NA06984, NA11840, NA06986, NA06989, NA06994, NA07346, NA07357, NA10851, NA11829, and NA11832.

The RNAseq libraries from *D. mojavensis* and *D. arizonae* are available through the NCBI Sequence Read Archive (SRA: http://www.ncbi.nlm.nih.gov/sra) under the accession no. SRX1272419 and SRX1277353.

The *A. tabida* dataset will be uploaded in SRA in the coming weeks.

## 3 RESULTS

### 3.1 Validation of the SNP calling method using available data from a model species

#### 3.1.1 Identification of variants

In order to evaluate the performance of our method, we needed to test it in the case where we knew which SNPs should be found. We thus focused on a dataset from humans in which SNPs were already annotated. We selected 2 populations (Toscans and Central Europeans) from the Geuvadis project [Lappalainen et al., 2013a], and downloaded the RNA-seq data of 10 individuals in each population. We sampled 10M reads from each individual and pooled individuals 5 by 5, to obtain two replicates of 5 pooled individuals per population. We ran KisSplice and Trinity on these 4 read sets and we aligned the variants of KisSplice to the Trinity transcripts using Blat (with at least 90% query coverage and 80% identity). Out of the 66530 bubbles initially found by KisSplice, 55590 (83%) mapped to Trinity-assembled transcripts, in 52345 different transcriptomic positions, 5797 partially aligned, and 5143 did not align. For our evaluation, we focused on the 52345 positioned bubbles.

To assess whether these bubbles were true SNPs, we first aligned the sequences of the variants (i.e. each path of the bubble) to the human reference genome and compared their genomic positions to a set of SNPs downloaded from the 1000 genome project webpage.

We also benchmarked our method against Gatk, a widely used method to call SNPs in the presence of a reference genome. Gatk was run with parameters recommended from the Gatk web page for RNA-seq data.

For each method, we calculated the Precision, i.e., the number of true SNPs out of the total number of predicted SNPs, and the Recall, i.e., the number of predicted SNPs out of the total number of true SNPs. Figure 5 summarises the overlap between the two methods and the set of true SNPs. First, we can observe that only a fraction of the true SNPs are recovered by either method. This result is expected because the set of true SNPs has been called using DNA-seq data, while the two methods benchmarked here only used RNA-seq data. Although we restricted our set of true SNPs to SNPs located in transcribed regions (at least 5 reads should cover the SNP), SNPs located in poorly expressed regions were harder to call. Hence, the recall of both methods is quite low: 37% for Gatk and 13% for KisSplice.

**Figure 5:**
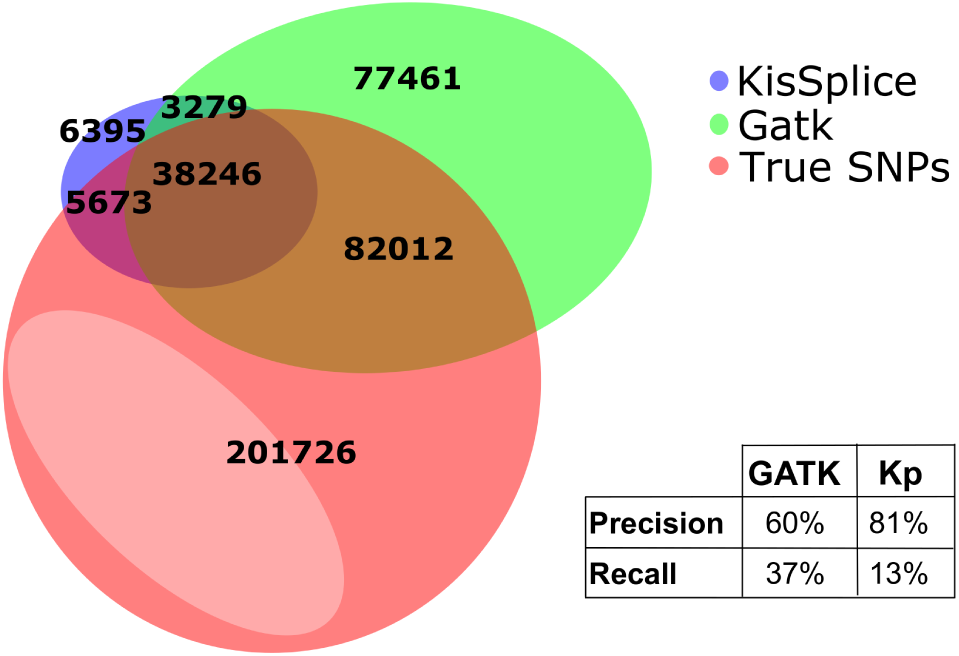
Precision and recall of KisSplice and Gatk. Half of the SNPs found by no method are covered by less than 10 reads (in light red).

Second, the number of SNPs predicted by Gatk was larger than the one predicted by KisSplice. This tendency is expected as mapping-based methods require fewer reads than assembly-based methods to call SNPs. However, Gatk tended to over-predict SNPs, with a precision of 60% while the precision of KisSplice was 81%. Both methods are comparable when we summarise these two results using the Balanced Accuracy measure (mean of Precision and Recall), which reached 48.5% for Gatk and 47% for KisSplice. **Importantly, KisSplice has the clear advantage that it does not require any reference genome, and can therefore be used for both model and non-model species**.

The low recall of both methods can be further discussed. A thorough inspection of the missed SNPs reveals that they were mostly located in poorly expressed genes and/or corresponded to rare alleles. Indeed, when the analysis was restricted to regions covered by at least 40 reads, both methods reached a recall of 70% (Supplementary Figure 3).

The default of precision can also be further analysed. We found that the false positives of Gatk were dominated by sequencing errors and mapping artifacts while the false positives of KisSplice were dominated by inexact repeats. In both cases, the impact of RNA editing was minor (less than 5% of cases were annotated in RADAR v2 [Ramaswami and Li, 2014]).

#### 3.1.2 Quantification of variants and statistical differential analysis

The quantification we obtain for variants called from pooled RNA-seq data reflects both the allele frequency of the variant in the pool and the expression level of the gene. An “expressed” allele frequency can be derived from these counts, by simply taking the ratio, but the obtained frequency is expected to be distorted compared to the allele frequency estimated from DNA-seq data. Several causes may be listed. First, within a heterozygous individual, one allele may be more expressed than the other, a process known as Allele Specific Expression (ASE). Second, RNA expression from different individuals (hence possibly different genotypes) can be variable within a pool, thus distorting the allelotype. In order to evaluate the magnitude of this distortion, we computed within each pool the correlations between the true allelic frequencies, and the estimated allele frequencies. To obtain the true allelic frequency within a pool, we took advantage of the availability of the genotypes of each individual from the Geuvadis dataset, and we simply summed up the number of alternative alleles over the total number of alleles within the pool. The expressed allele frequencies were obtained from KisSplice calls, summing the alternative allele counts of each individual over all allele counts of the pool.

We found that the distortion highly depends on the expression levels (Supplementary Figure 4). While the correlation was weak (0.65) for poorly expressed loci (less than 3 reads), it increased steadily with the expression level up to a plateau of 0.98. When we restricted to loci with at least 10 reads, the correlation reached 0.95. Similar results were obtained for every other pool and when we mixed up all 20 individuals within one pool (see Supplementary Figure 5). We therefore conclude that, whenever a locus was sufficiently expressed (at least 10 reads), the expressed allele frequency was a good predictor of the true allele frequency.

If we now compute the difference of allele frequencies across conditions (denoted by *df*), and compare it to the difference of expressed allele frequencies across conditions (denoted by *dfe*), the correlations are still high, but weaker, reaching a plateau of 80% for highly expressed loci (Supplementary Figure 6). The reason is that most SNPs do not have a significant difference of allele frequencies across our two populations, hence these correlations are contaminated by SNPs with (almost) equal allele frequencies. In this case, the difference of allele frequencies is just a random fluctuation. When considering all SNPs, the correlation between *df* and *dfe* is significant but weak (Figure 6-A)

**Figure 6:**
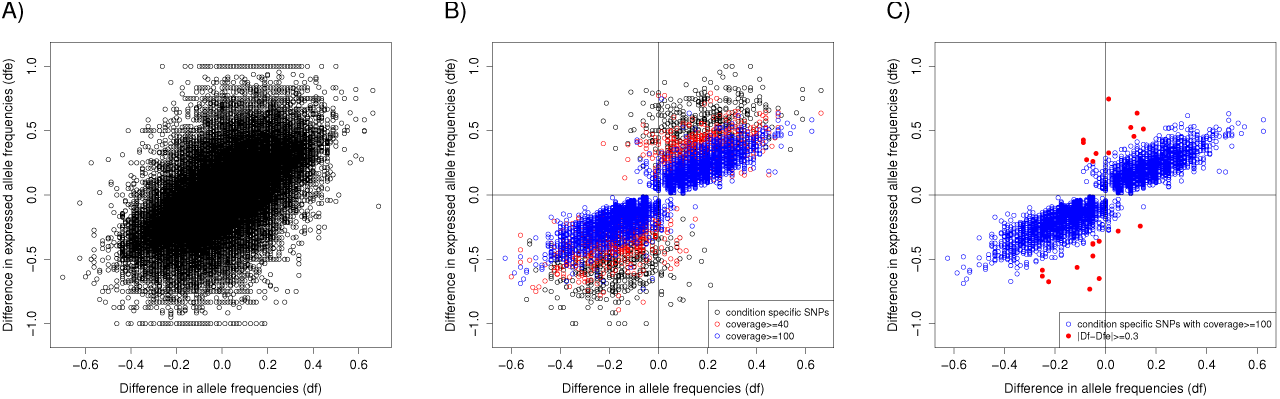
Difference of allele frequencies (df) Vs Difference of expressed allele frequencies (dfe) A)All SNPs B) Condition Specific SNPs C) Conditions Specific SNPs covered by at least 100 reads.

If we restrict to SNPs that are found as condition specific by KISSDE, then the correlation is much stronger (Figure 6-B). Finally, if we restrict to SNPs covered by a total of at least 100 reads (an average of 25 reads per sample), then the correlation is much higher (Figure 6-C). The more a gene is expressed, the higher the fit between *df* and *dfe*. A few SNPs (*n*=22) however exhibited a large difference between *df* and dfe (> 0.3). A detailed analysis of these cases reveals that they are located in immune genes (*n* = 5), in genes showing a very variable expression across individuals (*n* = 9), or in genes exhibiting an allele specific expression (*n* = 8).

**Overall, we conclude that, provided we restrict to condition specific SNPs, the metric we output with kissDE for the difference of expressed allele frequencies, that is** *dfe***, can largely be interpreted as a measure of the true difference of allele frequencies**.

#### 3.1.3 Functional impact

When no reference genome is available, it is not possible to obtain a genomic location for each SNP and therefore to apply SNPeff [Cingolani et al., 2012], or PolyPhen [Adzhubei et al., 2010], which are widely used softwares for assessing the impact of a SNP on the protein. In absence of any reference genome, a reference transcriptome can nevertheless be obtained, using a full-length transcriptome assembler like Trinity [Grabherr et al., 2011]. Based on this transcriptome, it is possible to assess the coding potential of each transcript using TransDecoder, to position the predicted SNPs onto the assembled transcripts using Blat [Kent, 2002], and finally to assess the impact of each SNP on its transcript(s). In the end, each positioned SNP is classified as coding or non coding. In the case where the SNP is located in the coding region, it is then classified as synonymous or non synonymous (See Methods).

Out of 52345 positioned SNPs (those which aligned to Trinity transcripts), 15582 cases (29%) fell in the CDSs and the other 71% fell in non-coding regions (including UTRs). Among the ones falling in the CDSs, we found that 51% (8063) were synonymous, while the other half (7519) were non synonymous.

To validate our predictions, we then intersected the genomic positions of our predicted SNPs with the genomic positions of SNPs in dbSNP, for which the functional impact is known. Out of the 52345 SNPs we predict, 41252 could be assigned a genomic position which matched a SNP annotated in dbSNP. Out of those 41252 cases, 39615 had a correct functional prediction. A thorough examination of the 1637 cases wrongly predicted reveals that in most cases, the transcript predicted by Trinity was very partial and was overlapping an intron (this happens when pre-mRNA is sampled together with mRNA at the RNA extraction step, despite selection of polyA+RNAs). In this case, the ORF predictor can over-predict coding regions, and our pipeline therefore tends to over-predict non synonymous cases. Figure 7 summarises our results for the prediction of functional impact. Overall, when SNPs can be evaluated, the precision of K2rt is 96%.

**Figure 7:**
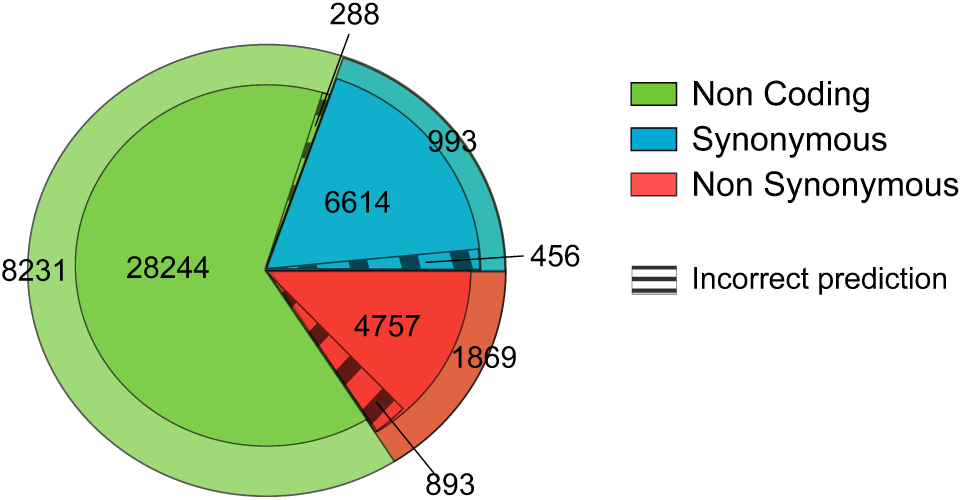
Results of KisSplice2RefTranscriptome. The green/red/blue areas correspond to Non Coding, Synonymous and Non Synonymous SNPs. The dashed area corresponds to errors of our functional predictions. The outer area corresponds to SNPs that are not in dbSNP and for which the functional prediction cannot be evaluated.

#### 3.1.4 Performance of the full pipeline

In the previous section, we evaluated our functional prediction independently of the remaining of our pipeline. We now turn to its evaluation within the full pipeline. Two situations can be discussed here. First, if only one experimental condition is considered, then no differential analysis is carried out. SNPs are identified and their functional impact is predicted. In this case, the prediction inherits from the errors made at the identification step. Out of 52345 predicted SNPs, 41252 are in dbSNP and 39615 have a functional impact correctly predicted. In the worst case scenario, if we consider that the 11093 SNPs for which there is no dbSNP entry are false positives, the precision of the pipeline is 77%. In practice, dbSNP is not exhaustive, and the true precision is between 77% and 96%. Second, if two conditions are considered (which is the original purpose of this study), then many of the false positives of the identification step are filtered out. Out of the 52345 predicted SNPs, 6033 are condition-specific, and 5507 have a correct functional prediction. Hence the precision for condition-specific SNPs is 91%.

### 3.2 Application of the method using biological data from species without any reference genome

From our study on the human dataset, we conclude that our method has a precision and recall similar to methods which require a reference genome. We now turn to the application of our method to non-model species.

#### 3.2.1 Application to Intraspecific Polymorphism: the case of *Asobara tabida*

We first applied our method to *Asobara tabida*, for which RNA-seq data from two lineages (SFR2 and SFR3) were available. These lineages come from the same population, but they differ by their phenotype of dependence to their symbiont *Wolbachia*. In the absence of *Wolbachia*, SFR2 individuals produce no eggs, while SFR3 produce some. Consequently, we suspect a low but significant genetic differentiation between lineages that could be associated with the phenotypes, or to genetic drift associated with maintenance in the laboratory. While the experimental design, with a single lineage for each phenotype, does not enable us to separate between these two effects, we think that this dataset is still well tailored for a validation of our method because: (a) no reference genome is available for this species; (b) individuals were pooled for RNA extraction; and (c) replicates are available for each lineage.

The transcriptomes of two replicates of pools of 30 individuals were sequenced through RNA-seq for each lineage, leading to 15M reads for each replicate. We ran our pipeline and found a total of 18609 positioned SNPs out of which 17031 are condition-specific. The large proportion of condition-specific SNPs is largely due to the fact that most of them are fixed in at least one lineage. Indeed, 21% of them are fixed in both lineages, 63% are fixed in one lineage and polymorphic in the other and 7% are polymorphic in both lineages (Supplementary Figure 7).

Out of the 17031 condition-specific variants, we found that 5608 (32%) were non coding, 6137 (36%) were synonymous and 3876 (22%) were non-synonymous.

Based on these results, we selected 27 cases for experimental validation: 10 were cases where the two lineages were fixed for a different nucleotide, 15 were cases where one lineage was fixed and the other polymorphic, 2 were cases where the two lineages were polymorphic. For all the 10 first cases, we were able to validate that the SNP was real and that the two lineages were indeed fixed for a different nucleotide (Figure 8). Out of the 17 remaining cases, we were able to validate that the SNP was real in all cases, but only in 9 cases were we able to validate that the site was polymorphic in one lineage (Figure 8). The rate of validation of the polymorphic status of the site within a lineage largely depended on the frequency of the minor allele (Supplementary Figure 8). Rare variants were harder to validate in terms of polymorphism detection. These rare variants could be false positives of our method, but they may also very well be true variants, not detectable experimentally using a direct sequencing technique without cloning. Importantly, although we could not always validate the fact that one site is polymorphic within a lineage, we systematically confirmed that the SNP was real, and that each lineage had a specific major allele. Therefore, we validated the condition-specificity of all SNPs.

**Figure 8:**
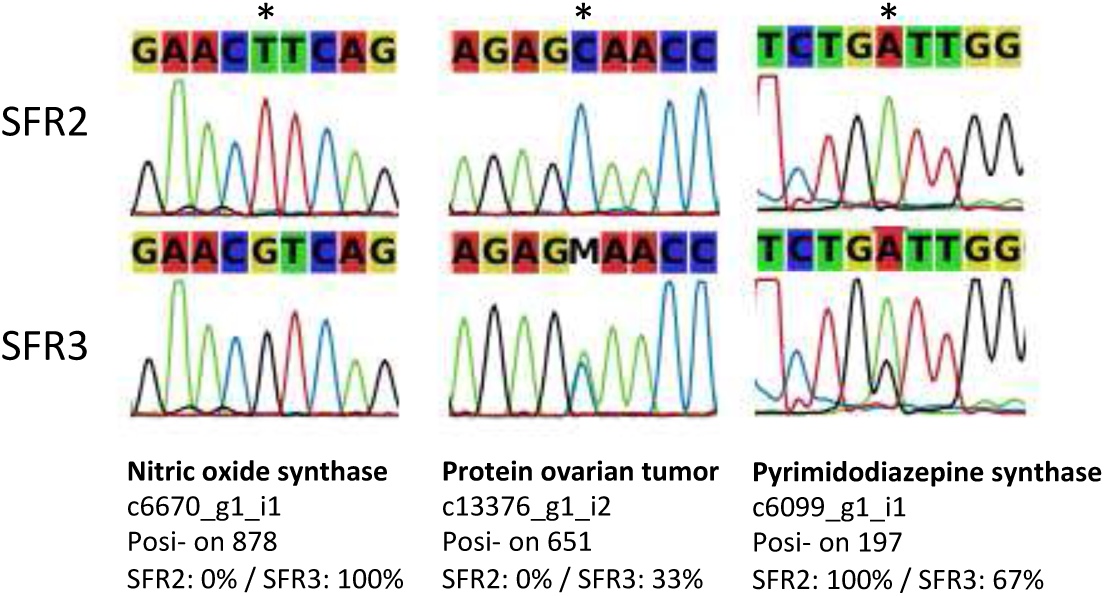
SNPs validated by Sanger sequencing. The first is fixed in both SFR2 and SFR3 lineages. The second and the third are polymorphic in SFR3

Because our RNA-seq data were initially obtained to compare the transcriptome of these two lineages, the design was not optimised for QTL analysis. In particular, each phenotype is represented by a single inbred genotype, making it difficult to separate the SNPs linked to the phenotype from those linked to drift. Despite this issue, we further characterised the functional impact of the condition-specific SNPs. Among all these genes, some called our attention regarding their possible implication in the dependence phenotype. For instance, some genes, such as *Dorsal* and *Hypoxia up-regulated protein 1*, presented SNPs in their UTRs and were differentially expressed between lineages. These genes are involved in immunity and oxidative stress homeostasis, two functions that have been shown as particularly important in this biological system. Another example concerns genes involved in oogenesis, like *OTU-domain containing protein* or *Female sterile*, that exhibit non-synonymous SNPs in their CDS regions. These few examples show how the suite we propose in this paper rapidly allows to link the SNPs detected to their functional impact, thus permitting to pinpoint candidate genes involved in phenotypic variation. Validation of these genes could involve either genetic studies *(e.g*., knock-down experiments) and/or other linkage analyses targeted to these candidates.

#### 3.2.2 Application to Interspecific Divergence: the case of *Drosophila mojavensis* and *Drosophila arizonae*

Similarly to the *Asobara* dataset, the drosophila dataset corresponds to non-model species, where individuals had to be pooled prior to RNA sequencing. In this case however, the two modalities of the phenotypes are not two populations of the same species, but two recently diverged species. This therefore enabled us to assess if our method also applies to a very different evolutionary scale, where differences of one nucleotide are no longer SNPs, but divergences. Additionally, the availability of the reference genome for *D. mojavensis* (and not *D. arizonae*) enabled us to study in depth the case of condition-specific inexact repeats.

*D. mojavensis* and *D. arizonae* are two closely related species that diverged 1MYA. We sequenced through RNA-seq the ovarian transcriptomes of two replicates of pools of 30 individuals for each species. We obtained 55M paired-end reads per replicate. We ran our pipeline on the data and obtained 51730 positioned SNPs, and most of them (51135) were condition-specific.

The condition-specific SNPs were mostly in coding regions (60%, *i.e*., 40674 SNPs). We could classify 34382 of them as synonymous, and the other 6292 SNPs as non-synonymous.

We selected 11 cases for experimental validation, 6 of which were divergent sites, and 5 were cases where the site was polymorphic in one species and fixed in the other. We were able to validate that the variation was condition-specific for all the divergent sites, and for 4 cases out of 5 for the polymorphic cases. Additionally, for 2 cases out of these 4, we were able to amplify the two alleles in the species where the site was predicted to be polymorphic.

In most cases, an observed variation in the transcriptome is caused by the presence of two alleles at one locus. However, it is also possible that two mono-allelic loci, if they exhibit the same sequence except for one nt, generate a variation that resembles a SNP. In order to quantify this phenomenon, we explicitely selected in the results of KisSplice, the variations for which one path was mapping to one locus and the other path was mapping to another locus. This was only possible because we had at our disposal a draft genome of *D. mojavensis*. We selected explicitely cases where we knew that the variation we detected was potentially caused by two loci. There were only 224 cases like this, which is very few compared to the total number of variations detected. We however tested 3 of them experimentally, and we were able to validate all of them. These cases are not true SNPs, but they correspond to recent paralog genes where one copy is more expressed in *D. arizonae*, and the other copy is more expressed in *D. mojavensis*.

## 4 Conclusion and Perspectives

We present a method that can discover condition-specific SNPs from raw RNA-seq data. The individuals may be pooled, which decreases the costs of library preparation, while still enabling to allelotype and to find variants specific to one condition. As no reference genome is required, the range of applications of the method is very large. We first evaluated our method in human, where a reference genome is available and SNPs are extensively annotated. We show that our method has similar performances in terms of precision and recall, compared to Gatk, a widely used mapping-based approach. We then evaluated our method on two non-model species.

In both cases, we were able to call variants, to classify them, and to discuss their functional impact. We selected a fraction of them for experimental validation through RT-PCR + Sanger sequencing. In all cases, we were able to validate that the variant was condition-specific. However, when the locus was predicted to be polymorphic in one condition, we were able to validate the presence of the two alleles only in cases where the minor allele frequency was at least 15%.

This work is a first approach towards transcriptome-wide association studies in nonmodel species. The method can readily be applied to RNA-seq data from any species, whenever two phenotypes are clearly identified and the goal is to find candidates for their genetic bases. In the case of continuous phenotypes, like height, the statistical framework can be generalised to quantitative trait loci (QTL).

This work focuses on SNP identification and analysis and does not address the question of the experimental design of a transcriptome-wide association study. A systematic evaluation of the optimal design is beyond the scope of this paper, but we would like to provide here briefly some basic advice. First, in all the case studies presented here, we considered only two replicates, which is the minimum required by our method. We clearly advise that for a pre-determined cost, it is wiser to have a low coverage for each replicate, but to increase the number of replicates. Second, the type of replicates to choose is probably a more central issue. In the case of *Asobara*, we sequenced two biological replicates, but both replicates were derived from the same lineage. Having replicates when extracting RNA is useful, but not as useful as replicates at the line-establishment step. Only this type of replicate can allow to discriminate between SNPs in the original population and genetic drift in the lab. Finally, if pooling is envisioned, the number of individuals per pool should be as large as possible, especially for very polymorphic species. The larger the pool, the more representative of the population it is.

From the point of view of our method itself, there is of course also room for improvement. In particular, we found that, while easy SNPs are identified by all methods, a large amount of difficult SNPs are currently being overseen. This is the case of SNPs located in repeated regions of the genome, and that are notoriously difficult to annotate. SNPs located very close to each other are also challenging to annotate. Without a reference genome, we found that they are particularly difficult to tell apart from inexact repeats. Finally, SNPs located within very polymorphic regions of the genome, like immune genes, are also very challenging, even for mapping-based approaches. The use of a single reference genome is clearly limiting. De novo assembly methods are a promising direction for these, but still need to be optimised.

For future work, we see two lines of research, which could ultimately be combined. First, we could take advantage of the availability of long reads coming from third generation sequencing platforms (Pacbio, Minion). In principle, long reads have the potential to solve most of the issues we mentioned, but currently, the error rates are too high (10–15%) and the sequencing depth is not sufficient to apply to RNA-seq. In the meantime, it seems still relevant to keep on working in the context of short reads, but we think that the best resolution we can achieve for the prediction of difficult SNPs is not well captured by sequences. Graphs could instead well represent close SNPs and a partial quantification of their phasing.

## 6 Acknowledgments

The authors would like to thank Marie-Christine Charpentier and Delphine Charif for help on the analysis of the *Asobara* dataset; Gustavo Sacomoto for help on the analysis of the Geuvadis dataset; Sebastien Deraison for help on experimental validation of the candidates of Drosophila

## Footnotes

1 http://gatkforums.broadinstitute.org/discussion/3892/the-gatk-best-practices-for-variant-calling--on-rnaseq-in-full-detail posted on 2014–03–06, last updated on 2014–10–31

## References

Abecasis G. R., Auton A., Brooks L. D., DePristo M. A., Durbin R. M., Handsaker R. E., Kang H. M., Marth G. T., and McVean G. A. (2012). An integrated map of genetic variation from 1,092 human genomes. Nature, 491(7422):56–65.

Adzhubei I. A., Schmidt S., Peshkin L., Ramensky V. E., Gerasimova A., Bork P., Kondrashov A. S., and Sunyaev S. R. (2010). A method and server for predicting damaging missense mutations. Nature methods, 7(4):248–249.

Anders S. (2010). Analysing rna-seq data with the deseq package. Mol Biol, 43(4):1–17.

Bullard J. H., Purdom E., Hansen K. D., and Dudoit S. (2010). Evaluation of statistical methods for normalization and differential expression in mrna-seq experiments. BMC bioinformatics, 11(1):94.

Chikhi R. and Rizk G. (2013). Space-efficient and exact de Bruijn graph representation based on a Bloom filter. Algorithms for molecular biology: AMB, 8(1):22.

Cingolani P., Platts A., Wang L. L., Coon M., Nguyen T., Wang L., Land S. J., Lu X., and Ruden D. M. (2012). A program for annotating and predicting the effects of single nucleotide polymorphisms, snpeff: Snps in the genome of drosophila melanogaster strain w1118; iso-2; iso-3. Fly, 6(2):80–92.

Dedeine F., Vavre F., Fleury F., Loppin B., Hochberg M. E., and Bouletreau M. (2001). Removing symbiotic Wolbachia bacteria specifically inhibits oogenesis in a parasitic wasp. P. Natl. Acad. Sci. USA, 98(11):6247–52.

Dedeine F., Vavre F., Shoemaker D. D., and Boulétreau M. (2004). Intra-individual coexistence of a Wolbachia strain required for host oogenesis with two strains inducing cytoplasmic incompatibility in the wasp Asobara tabida. Evolution, 58(10):2167–74.

Dillies M.-A., Rau A., Aubert J., Hennequet-Antier C., Jeanmougin M., Servant N., Keime C., Marot G., Castel D., Estelle J., et al. (2013). A comprehensive evaluation of normalization methods for illumina high-throughput rna sequencing data analysis. Briefings in bioinformatics, 14(6):671–683.

Dobin A., Davis C. A., Schlesinger F., Drenkow J., Zaleski C., Jha S., Batut P., Chaisson M., and Gingeras T. R. (2013). Star: ultrafast universal rna-seq aligner. Bioinformatics, 29(1):15–21.

Engström P. G., Steijger T., Sipos B., Grant G. R., Kahles A., Alioto T., Behr J., Bertone P., Bohnert R., Campagna D., Davis C. a., Dobin A., Gingeras T. R., Goldman N., Guigó, R., Harrow J., Hubbard T. J., Jean G., Kosarev P., Li S., Liu J., Mason C. E., Molodtsov V., Ning Z., Ponstingl H., Prins J. F., Rätsch G., Ribeca P., Seledtsov I., Solovyev V., Valle G., Vitulo N., Wang K., Wu T. D., and Zeller G. (2013). Systematic evaluation of spliced alignment programs for RNA-seq data. Nature methods, 10(12).

Grabherr M. G., Haas B. J., Yassour M., Levin J. Z., Thompson D. a., Amit I., Adiconis X., Fan L., Raychowdhury R., Zeng Q., Chen Z., Mauceli E., Hacohen N., Gnirke A., Rhind N., di Palma F., Birren B. W., Nusbaum C., Lindblad-Toh K., Friedman N., and Regev A. (2011). Full-length transcriptome assembly from RNA-Seq data without a reference genome. Nature biotechnology, 29(7):644–52.

Iqbal Z., Caccamo M., Turner I., Flicek P., and McVean G. (2012). De novo assembly and genotyping of variants using colored de Bruijn graphs. Nature genetics, 44(2):226–32.

Kent W. J. (2002). Blatthe blast-like alignment tool. Genome research, 12(4):656–664.

Kremer N., Dedeine F., Charif D., Finet C., Allemand R., and Vavre F. (2010). Do variable compensatory mechanisms explain the polymorphism of the dependence phenotype in the Asobara tabida-wolbachia association? Evolution, 64(10):2969–2979.

Kremer N., Voronin D., Charif D., Mavingui P., Mollereau B., and Vavre F. (2009). Wolbachia interferes with ferritin expression and iron metabolism in insects. PLoS Pathog., 5(10):e1000630.

Lappalainen T., Sammeth M., Friedländer M. R., ACt Hoen P., Monlong J., Rivas M. A., Gonzalez-Porta M., Kurbatova N., Griebel T., Ferreira P. G., et al. (2013a). Transcriptome and genome sequencing uncovers functional variation in humans. Nature, 501(7468):506–511.

Lappalainen T., Sammeth M., Friedländer M. R., ’t Hoen, P. A. C., Monlong J., Rivas M. A., Gonzàlez-Porta M., Kurbatova N., Griebel T., Ferreira P. G., Barann M., Wieland T., Greger L., van Iterson M., Almlöf J., Ribeca P., Pulyakhina I., Esser D., Giger T., Tikhonov A., Sultan M., Bertier G., MacArthur D. G., Lek M., Lizano E., Buermans, H. P. J., Padioleau I., Schwarzmayr T., Karlberg O., Ongen H., Kilpinen H., Beltran S., Gut M., Kahlem K., Amstislavskiy V., Stegle O., Pirinen M., Montgomery S. B., Donnelly P., McCarthy M. I., Flicek P., Strom T. M., Lehrach H., Schreiber S., Sudbrak R., Carracedo A., Antonarakis S. E., Hasler R., Syvänen A.-C., van Ommen G.-J., Brazma A., Meitinger T., Rosenstiel P., Guigó, R., Gut I. G., Estivill X., and ermitzakis E. T. (2013b). Transcriptome and genome sequencing uncovers functional variation in humans. Nature, 501(7468):506–11.

Leggett R. M., Ramirez-Gonzalez R. H., Verweij W., Kawashima C. G., Iqbal Z., Jones, J. D. G., Caccamo M., and Maclean D. (2013). Identifying and classifying trait linked polymorphisms in non-reference species by walking coloured de bruijn graphs. PloS one, 8(3):e60058.

Matzkin L. M. (2004). Population genetics and geographic variation of alcohol dehydrogenase (adh) paralogs and glucose-6-phosphate dehydrogenase (g6pd) in drosophila mojavensis. Molecular Biology and Evolution, 21(2):276–285.

Modolo L. and Lerat E. (2015). Urqt: an efficient software for the unsupervised quality trimming of ngs data. BMC bioinformatics, 16(1): 137.

Peterlongo P., Schnel N., Pisanti N., Sagot M. F., and Lacroix V. (2010). Identifying SNPs without a reference genome by comparing raw reads. Lecture Notes in Computer Science (including subseries Lecture Notes in Artificial Intelligence and Lecture Notes in Bioinformatics), 6393 LNCS:147–158.

Pevzner P. a., Tang H., and Waterman M. S. (2001). An Eulerian path approach to DNA fragment assembly. Proceedings of the National Academy of Sciences of the United States of America, 98(17):9748–9753.

Piskol R., Ramaswami G., and Li J. B. (2013). Reliable identification of genomic variants from RNA-seq data. American Journal of Human Genetics, 93(4): 641–651.

Ramaswami G. and Li J. B. (2014). RADAR: a rigorously annotated database of A-to-I RNA editing. Nucleic Acids Research, 42(D1):D109–D113.

Reed L., Nyboer M., and Markow T. (2007). Evolutionary relationships of drosophila mojavensis geographic host races and their sister species drosophila arizonae. Molecular Ecology, 16(5):1007–1022.

Romiguier J., Gayral P., Ballenghien M., Bernard, a., Cahais V., Chenuil, a., Chiari Y., Dernat R., Duret L., Faivre N., Loire E., Lourenco J. M., Nabholz B., Roux C., Tsagkogeorga G., a. T. Weber, a., Weinert L. a., Belkhir K., Bierne N., Glémin S., and Galtier N. (2014). Comparative population genomics in animals uncovers the determinants of genetic diversity. Nature, (V).

Sacomoto G., Sinaimeri B., Marchet C., Miele V., Sagot M.-F., and Lacroix V. (2014). Navigating in a Sea of Repeats in RNA-seq without Drowning. Lecture Notes in Bioinformatics, 8701: 82–96.

Sacomoto G. a. T., Kielbassa J., Chikhi R., Uricaru R., Antoniou P., Sagot M.-F., Peterlongo P., and Lacroix V. (2012). KISSPLICE: de-novo calling alternative splicing events from RNA-seq data. BMC bioinformatics, 13 Suppl 6(Suppl 6):S5.

Salikhov K., Sacomoto G., and Kucherov G. (2014). Using cascading Bloom filters to improve the memory usage for de Brujin graphs. Algorithms for Molecular Biology, 9(1):2.

Schlötterer C., Tobler R., Kofler R., and Nolte V. (2014). Sequencing pools of individuals mining genome-wide polymorphism data without big funding. Nature Reviews Genetics, 15(11):749–763.

Tilgner H., Knowles D. G., Johnson R., Davis C. A., Chakrabortty S., Djebali S., Curado J., Snyder M., Gingeras T. R., and Guigó, R. (2012). Deep sequencing of subcellular RNA fractions shows splicing to be predominantly co-transcriptional in the human genome but inefficient for lncRNAs. Genome research, 22(9):1616–25.

Uricaru R., Rizk G., Lacroix V., Quillery E., Plantard O., Chikhi R., Lemaitre C., and Peterlongo P. (2015). Reference-free detection of isolated SNPs. Nucleic acids research, 43(2):e11.

Zerbino D. R. and Birney E. (2008). Velvet: algorithms for de novo short read assembly using de Bruijn graphs. Genome research, 18(5):821–9.

